# Formulation, Evaluation and Antibacterial Efficiency of water-based herbal Hand Sanitizer Gel

**DOI:** 10.1101/373928

**Authors:** Shri Balakrishna Acharya, Saradindu Ghosh, Giriraj Yadav, Kavita Sharma, Sirsendu Ghosh, Sushil Joshi

## Abstract

Hands are the primary mode of transmission of microbes and infections. Hand-washing is critical in food production, food service and also important in healthcare setting, homes and day care preparations. The present research was aimed to evaluate the antibacterial efficacy of various herbal oils such as Cinnamon oil, Eucalyptus oil, menthol oil and lavender oil and found that cinnamon oil showed better antibacterial activity. Also the research was carried out to formulate and evaluate the poly herbal Hand wash gel containing *Azadirachta indica, Ocimum sanctum* and *Citrus limon* extracts. The anti-microbial activity of the formulated herbal hand wash gel was tested against *Escherichia coli, Staphylococcus aureus* and *Salmonella* by spread plate techniques and the results obtained were compared with commercial antibacterial standards. Also the efficiency was checked by using the hand wash gel on volunteers. The results from the present work suggest and support the incorporation and utilization of herbs in the formulations to give better effect.

## Introduction

Hands are primary mode of transmission of microbes and infections, Hand hygiene is therefore the most important measure to avoid the transmission of harmful germs and prevent the infections. Hand hygiene is the single most important, simplest, and least expensive means of preventing nosocomial infections. [1] Contaminated hands can serve as vectors for the transmission of microorganisms. Pathogenic microorganisms accountable for outbreaks are spread from the hands of the food handler to others when the food handler contaminates his/her hands and then passes these microorganisms to consumers via hand contact with food or drinks. The consumer is exposed following the ingestion of these microorganisms, which may cause gastrointestinal illness. Hand contact with ready-to-eat foods represents a very important mechanism by which pathogens may enter the food supply. Food handlers whose work involves touching unwrapped foods to be consumed raw or without further cooking or other forms of treatment have been identified as a particular risk group [2].

To protect the skin from harmful micro organisms and to prevent spreading of many contagious diseases, hand washing is absolutely an important precaution. Food production workers and foodservice personnel must be taught to use correct hand and fingertip washing by management in preparation for work [3]. Any health-care worker, caregiver or person involved in direct or indirect patient care needs to be concerned about hand hygiene and should be able to perform it correctly and at the right time [4]. Black et al. (1981) reported a study that demonstrated a decline in diarrheal illnesses (due to Shigella, Giardia and rotavirus) in day care centers when employees were taught to use good hand washing procedures [5]. Hand washing removes visible dirt from hands and reduce the number of harmful microorganisms such as, E. coli and Salmonella can be carried by people, animals or equipment and transmitted to food [6]. By far the most common mode of pathogen transmission to food by the infected food handler is via faecally contaminated hands. Poor hand hygiene is the contributing factor [2]. WHO has recommended all people should wash hands before during and after preparing food, before eating food, before and after caring for someone who is sick, before and after treating a cut or wound, after using the toilet and changing diapers or cleaning up a child who has used the toilet. After blowing your nose, coughing, or sneezing, after touching an animal or animal waste, after handling pet food or pet treats and after touching garbage [7]. For generations, hand washing with soap and water has been considered a measure of personal hygiene. The concept of cleansing hands with an antiseptic agent probably emerged in the early 19th century. As early as 1822, a French pharmacist demonstrated that solutions containing chlorides of lime or soda could eradicate the foul odors associated with human corpses and that such solutions could be used as disinfectants and antiseptics. In a paper published in 1825, this pharmacist stated that physicians and other persons attending patients with contagious diseases would benefit from moistening their hands with a liquid chloride solution [8].

Several studies suggested that, sanitizers with at least 70% alcohol were suggested to kill 99.9% of the bacteria on hands [13]. Alcohol-based hand sanitizers exist in liquid, foam, and easy-flowing gel formulations. Sometimes combined with quats (quaternary ammonium cations) such as benzalconium chloride quarts are added at level up to 200parts per million to increase antimicrobial effectiveness [14].

Before the discovery of modern medicine, plants were the main remedy for treating various diseases. With the advent of different antibiotics microbes also gradually develop resistance to these substances. These bring researchers interest towards the plants having antimicrobial properties. They try to exploit the unique ability of different secondary metabolites to show persistent and prolonged activity against broad spectrum of microbes [15].

In this study we used *Azadirachta indica, Ocimum sanctum* and *Citrus limon* due to their individual benefits. All parts of *Azadirachta indica* are used in many medicinal treatments like skin diseases, healthy hair, improve liver function; detoxify the blood, anti-inflammatory, anti-diabetic, antiviral, ant carcinogenic, immune-modulatory etc. Aqueous extract of stem bark has been shown to enhance the immune response of Balb-c mice to sheep red blood cells in –vivo [16]. *Ocimum sanctum* is an aromatic plant in the family Lamiaceae which is native to the Indian subcontinent. Ayurvedic system of medicine of *Ocimum sanctum* for various aliment and has been show to significant anti-stress property. Different parts of the plant to be effectively in a number of diseases [16]. *Citrus limon* is an important medicinal plant of the family Rutaceae. It is cultivated mainly for its alkaloids, which are having antibacterial potential in crude extracts of different parts (viz, leaves, stem, root and flower) of lemon against clinically significant bacterial strains has been reported [17]. Citrus flavanoid has a large spectrum of biological activity including antibacterial, antifungal anti-diabetic, anticancer and antiviral activities [18].

Traditionally, microbes incorporating hands are divided into resistant and transient flora resistant flora E.g. Staphylococcus azures and staphylococcus epidermis which always canonizing deeper skin layers which always resistant to mechanical removing and had over pathogenic potential Transient flora E.g. *Staphylococcus aureus* are the basically colonizes the superficial skin layers for short periods [19]. Alcohol kills germs by destroying the cell membranes and denaturing proteins of bacterial cells. Because of this gram negative bacteria (e.g. *E.coli* and *Salmonella)* are more susceptible to sanitizers, science they have a thin peptidoglycan cell wall surrounded by an outer membrane, which can be dissolved by alcohols. Gram positive bacteria (Staphylococcus) have a thicker peptidoglycan cell wall and are less vulnerable to alcohol based sanitizers, killed by these sanitizers there are some non alcohol based sanitizers that are more effective against gram positive bacteria, such as benzalconium chloride (BAC) based sanitizer.

## MATERIALS AND METHODS

The study was conducted between January 2017 and June 2017. The study was carried out in the Department of Microbiology, Patanjali Natural Coloroma Pvt Ltd.

### Chemicals and Regents

The *Azadirachta indica, Ocimum sanctum* and *Citrus limon* were collected from Patanjali Natural Coloroma Pvt. Ltd. Haridwar, Uttrakhand, India. All other reagents/chemicals were used as analytical grade.

### Extraction

After sun drying, the material is made into powder with the help of grinder and each plant material is weighed. 50 gm of coarse powder of plant material to be extracted separately in 400 ml of water using sox let extractor for 6hrs/sample. Temperature to be maintained till the boiling point of water i.e. 100°c. The extract is separated from the solvent with the help of rotary evaporation. The solvent vaporizes, leaving the extract which is then oven-dried.

### Samples

A. **Volunteers Sample**: Swabs from hand skin of volunteers without any clinical signs of infection were included in his study.
B. **Bacterial samples**: Gram positive and Gram negative bacteria i.e. *Staphylococcus aureus, Escherichia coli, Salmonella sp* and *Candida albicans* were collected from IMTEC Chandigarh, India. Bacterial suspension of concentration 10CFU/ml was used.

### Media

A. Nutrient broth and agar
B. MacConkey agar
C. Mueller Hinton Agar

### Methods

#### Standardization of inoculums

The inoculums prepared from the stock cultures, were maintained on nutrient agar at 4°C and sub cultured onto Nutrient broth using a sterile wire loop.

#### Antimicrobial Studies of herbal extracts

The screening of anti-microbial efficacy of the herbal extracts was performed on various micro organisms by using dip well method as per standard procedure. Three sterile petri plates were taken for testing the antimicrobial activity of herbal extracts against three different microorganisms i.e. *Escherichia coli, Staphylococcus aureus, Salmonella* and *Candida albicans*. The plates were filled with MacConkey and Muller Hinton agar solution and allowed for solidification. After solidification the microorganisms from the subculture were inoculated into the nutrient agar media and three discs were inoculated with *Azadirachta indica, Ocimum sanctum* and *Citrus limon* extracts respectively. The plates were incubated at 37°C for overnight. After 24 hours of incubation, the plates were observed for the zone of inhibition. From the zone of inhibition the anti microbial activity of formulation is estimated as shown in the Table (1, 2, 3) below.

#### Determination of Minimum Inhibitory Concentration (MIC) of the extracts

The MIC is defined as the lowest concentration that completely inhibits the growth of microorganisms for 24 hrs incubation. Determination of minimum inhibitory concentration of extracts was determined by preparing different concentrations of extracts 200μg, 400μg and 800μg were added respectively to the nutrient broth (Table 1,2,3). A 50μl volume of each dilution was added aseptically into the wells of Mueller Hinton agar plates that were already seeded with the standardized inoculums of the test bacteria. All experiments were performed in triplicate. The agar plates were incubated at 37°C for 24 hours. The lowest concentration of extracts showing a clear zone of inhibition was considered as the MIC.

**Table 1:** Result of antimicrobial study of *Azadirachta indica* (mean ± SD) (n=3)

| Concn of Drug ($\mu\text{g/ml}$)/<br>*Organisms* | Zone of inhibition (diameter mm) | | | | |
| --- | --- | --- | --- | --- | --- |
|  | 800 | 400 | 200 | Control | STD |
| *E. coli* | 32 ± 2.0 | 30 ± 1.2 | 29 ± 1.5 | 0 | 24 ± 1.1 |
| *S. aureus* | 28 ± 1.8 | 27 ± 2.0 | 26 ± 1.9 | 0 | 27 ± 1.1 |
| *Salmonella Sp.* | 24 ± 1.2 | 16 ± 1.7 | 15 ± 1.5 | 0 | 19 ± 1.7 |
| *C. albicans* | 26 ± 1.6 | 20 ± 1.4 | 19 ± 1.5 | 0 | 28 ± 1.4 |

**Table 2:** Result of antimicrobial study of *Ocimum sanctum* (mean ± SD) (n=3)

| Concn of Drug ($\mu\text{g/ml}$)/<br>*Organisms* | Zone of inhibition (diameter mm) | | | | |
| --- | --- | --- | --- | --- | --- |
|  | 800 | 400 | 200 | Control | STD |
| *E. coli* | 28 ± 2.0 | 18 ± 1.2 | 15 ± 1.5 | 0 | 24 ± 1.1 |
| *S. aureus* | 25 ± 1.8 | 17 ± 2.0 | 16 ± 1.9 | 0 | 27 ± 1.1 |
| *Salmonella Sp.* | 20 ± 1.2 | 17 ± 1.7 | 15 ± 1.5 | 0 | 19 ± 1.7 |
| *C. albicans* | 28 ± 1.6 | 18 ± 1.4 | 14 ± 1.5 | 0 | 28 ± 1.4 |

**Table 3:** Result of antimicrobial study of *Citrus limon* (mean ± SD) (n=3)

| Concn of Drug ($\mu\text{g/ml}$)/<br>*Organisms* | Zone of inhibition (diameter mm) | | | | |
| --- | --- | --- | --- | --- | --- |
|  | 800 | 400 | 200 | Control | STD |
| *E. coli* | 30 ± 2.0 | 28 ± 1.2 | 21 ± 1.5 | 0 | 24 ± 1.1 |
| *S. aureus* | 34 ± 1.8 | 27 ± 2.0 | 26 ± 1.9 | 0 | 27 ± 1.1 |
| *Salmonella Sp.* | 32 ± 1.2 | 16 ± 1.7 | 15 ± 1.5 | 0 | 19 ± 1.7 |
| *C. albicans* | 36 ± 1.6 | 20 ± 1.4 | 19 ± 1.5 | 0 | 28 ± 1.4 |

#### Formulation of Herbal Hand Sanitizer Gel

Carbapol 940 and EDTA were added to deionized water with constant stirring. After uniform mixing, glycerine was added with slow stirring to avoid formation of possible air bubbles in the product. All the extracts were added to this mixture followed by the addition of PG. Finally, all the aqueous extract of *Azadirachta indica, Ocimum sanctum* and *Citrus limon* were added along with 0.3% of perfume and mixed by slow stirring to obtain uniform product. Prepared products were stored in HDPE containers.

### Characterization of herbal hand washes gel

#### pH

The pH was determined by using digital pH meter and the pH of herbal hand wash was found 6.5±0.1.

#### Viscosity

The viscosity of hand wash was determined by using digital Brookfield viscometer. Measured quantity of herbal hand wash was taken into a beaker and the tip of viscometer was immersed into the hand wash gel and the viscosity was measured in triplicate. The viscosity was found 50c Pascals.

#### Antimicrobial studies of herbal hand wash gel

The screening of anti-microbial efficacy of the formulated poly herbal hand wash gel was aseptically performed on *Escherichia coli, Staphylococcus aureus*, and *Salmonella* by using Dip well Agar Diffusion Technique described by Bauer et al., 19669 and demonstrated by Cakir et al., 200410 was employed for antibacterial bioassay. A well was prepared in the plates (containing 15ml of Muller– Hinton agar medium) with the help of a cork-borer (0.85cm). 100μl of the test compound (herbal hand wash gel) was introduced into the well. The standard antibiotic discs like erythromycin, penicillin, streptomycin and ampicillin were used as a standard. The plates were incubated overnight at 37°C. Efficiency of hand wash gel was determined by measuring the diameter of zone of inhibition at 200μg ml^−1^, 400μgml^−1^ and 800μgml^−1^ concentration. (Table 4, Fig 1)

**Fig 1:**
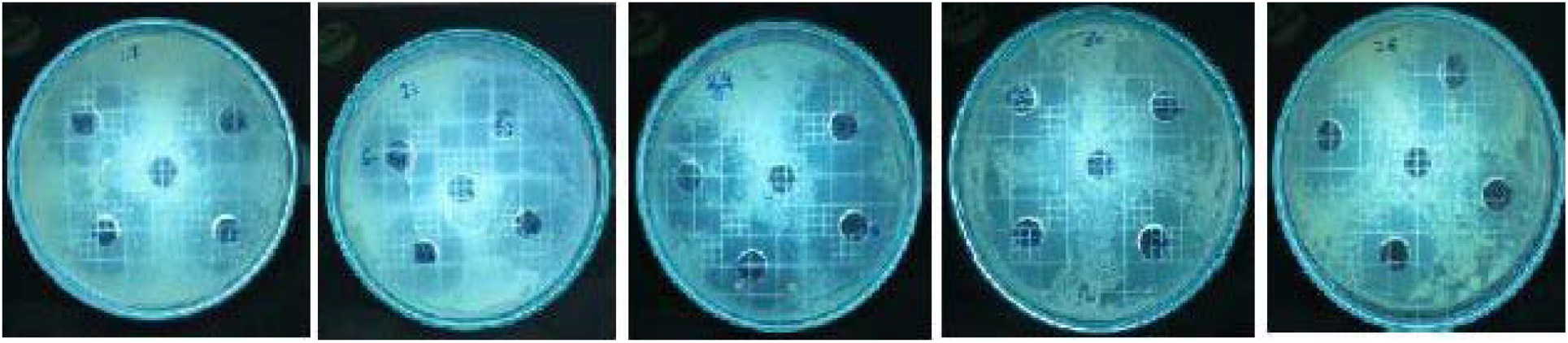
Antimicrobial activity of Hand sanitizer against various pathogens

**Table 4:** Result of antimicrobial study of Hand-sanitizer (mean ± SD) (n=3)

| Concn of Drug (µg/ml)/<br>*Organisms* | Zone of inhibition (diameter mm) |  |  |  |  |
| --- | --- | --- | --- | --- | --- |
|  | 800 | 400 | 200 | Control | STD |
| *E. coli* | 36 ± 2.0 | 32 ± 1.2 | 29 ± 1.5 | 0 | 32 ± 1.1 |
| *S. aureus* | 32 ± 1.8 | 30 ± 2.0 | 28 ± 1.9 | 0 | 31 ± 1.1 |
| *Salmonella Sp.* | 37 ± 1.2 | 30 ± 1.7 | 25 ± 1.5 | 0 | 20 ± 1.7 |
| *C. albicans* | 31 ± 1.6 | 28 ± 1.4 | 21 ± 1.5 | 0 | 28 ± 1.4 |

#### Antibacterial efficiency of herbal hand sanitizer gel on volunteers

The antibacterial efficiency was performed by spread plate technique. Samples were collected from the five different volunteers showing no clinical signs of dermal abrasion, trauma and infection. Approximately 500 μl of herbal hand wash gel was applied to both hands. After washing the hands, the samples were collected from each volunteer in a separate glass beaker and the collected samples were allowed to grow on nutrient agar media for overnight at 37□C and per ml CFU were calculated. (Fig 2, 3, 4) at 3 different randomly selected day.

**Fig 2:**
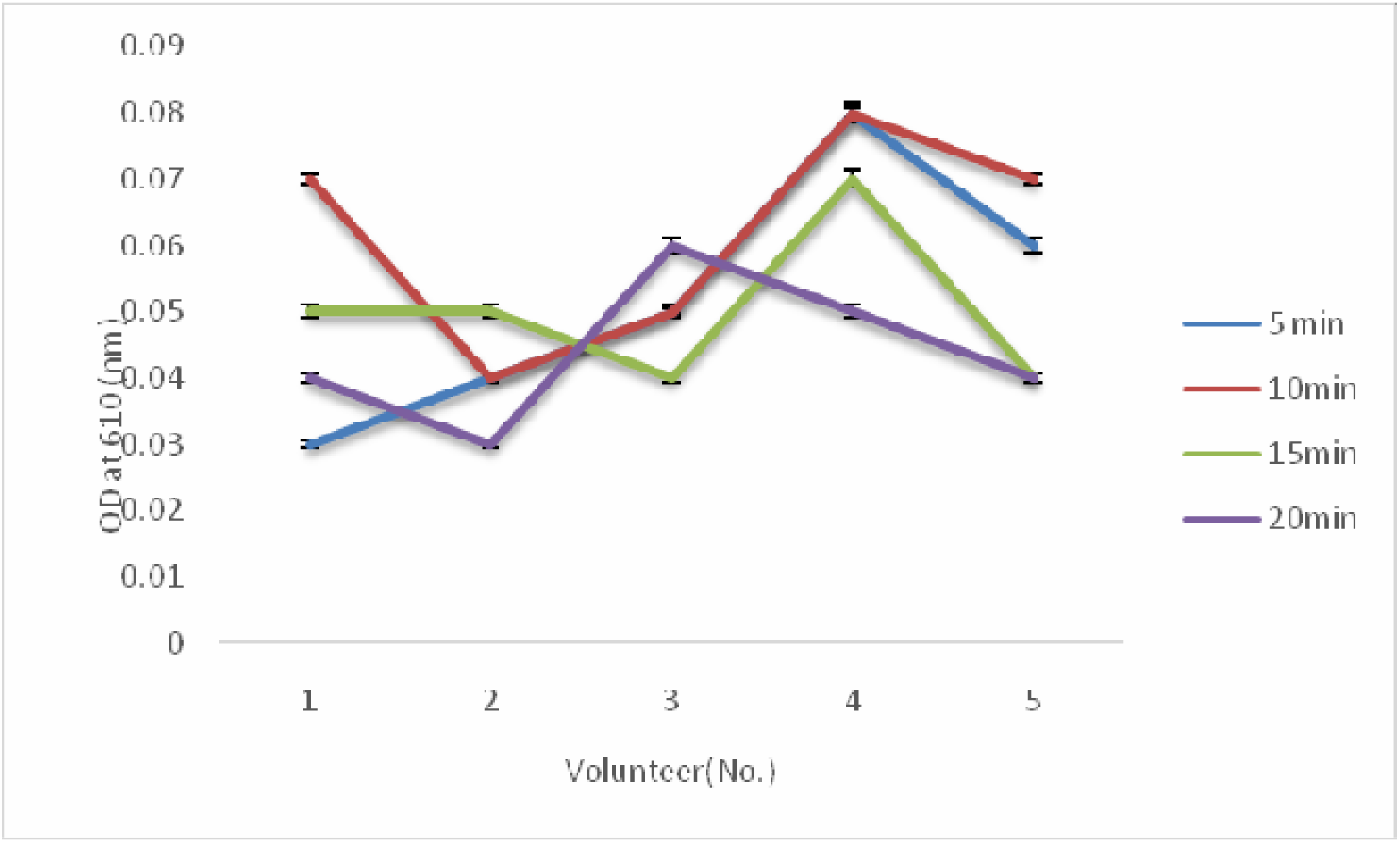
1st day observation of antimicrobial activity of Hand sanitizer against hand swab.

**Fig 3:**
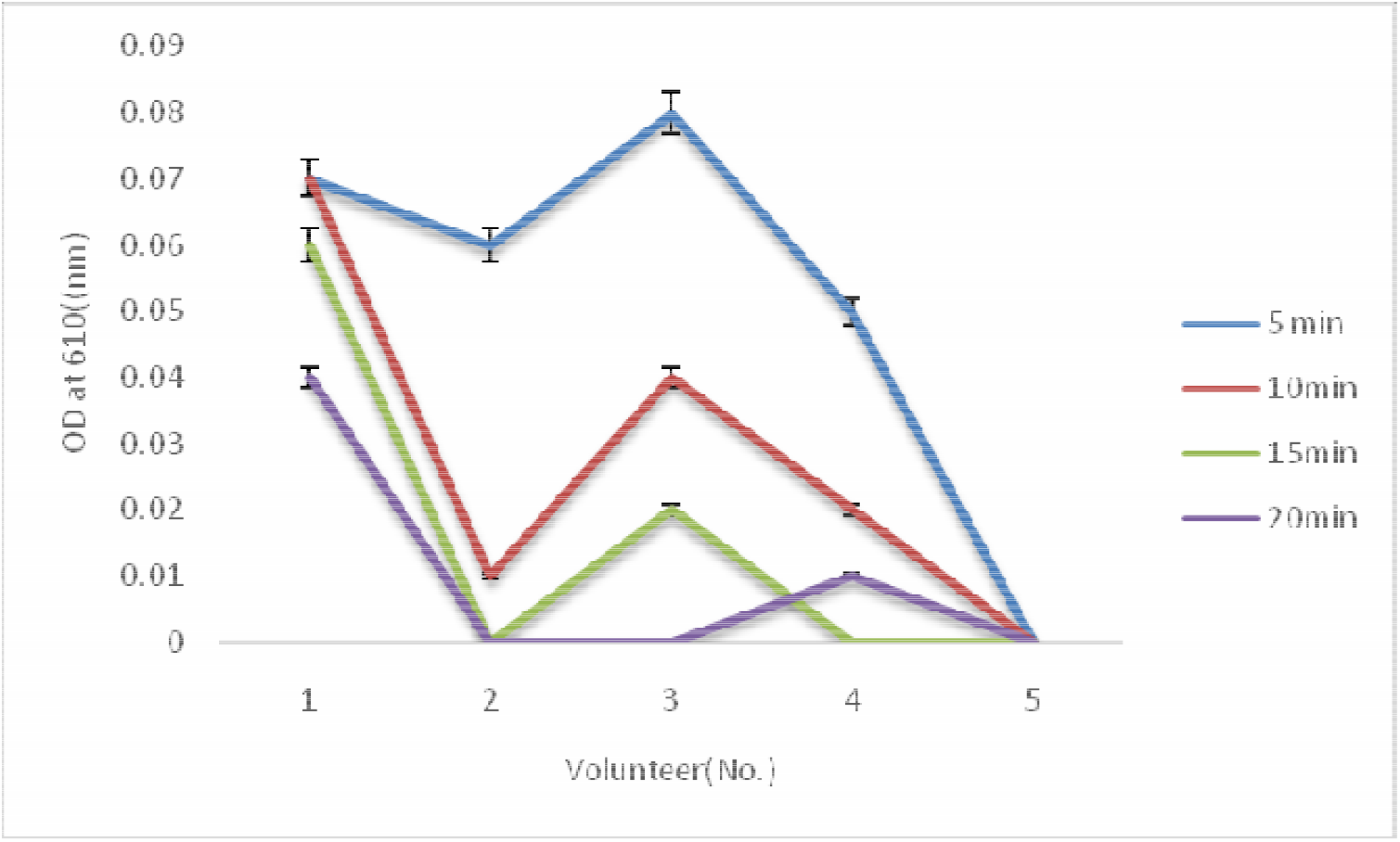
2nd day observation of antimicrobial activity of Hand sanitizer against hand swab

**Fig 4:**
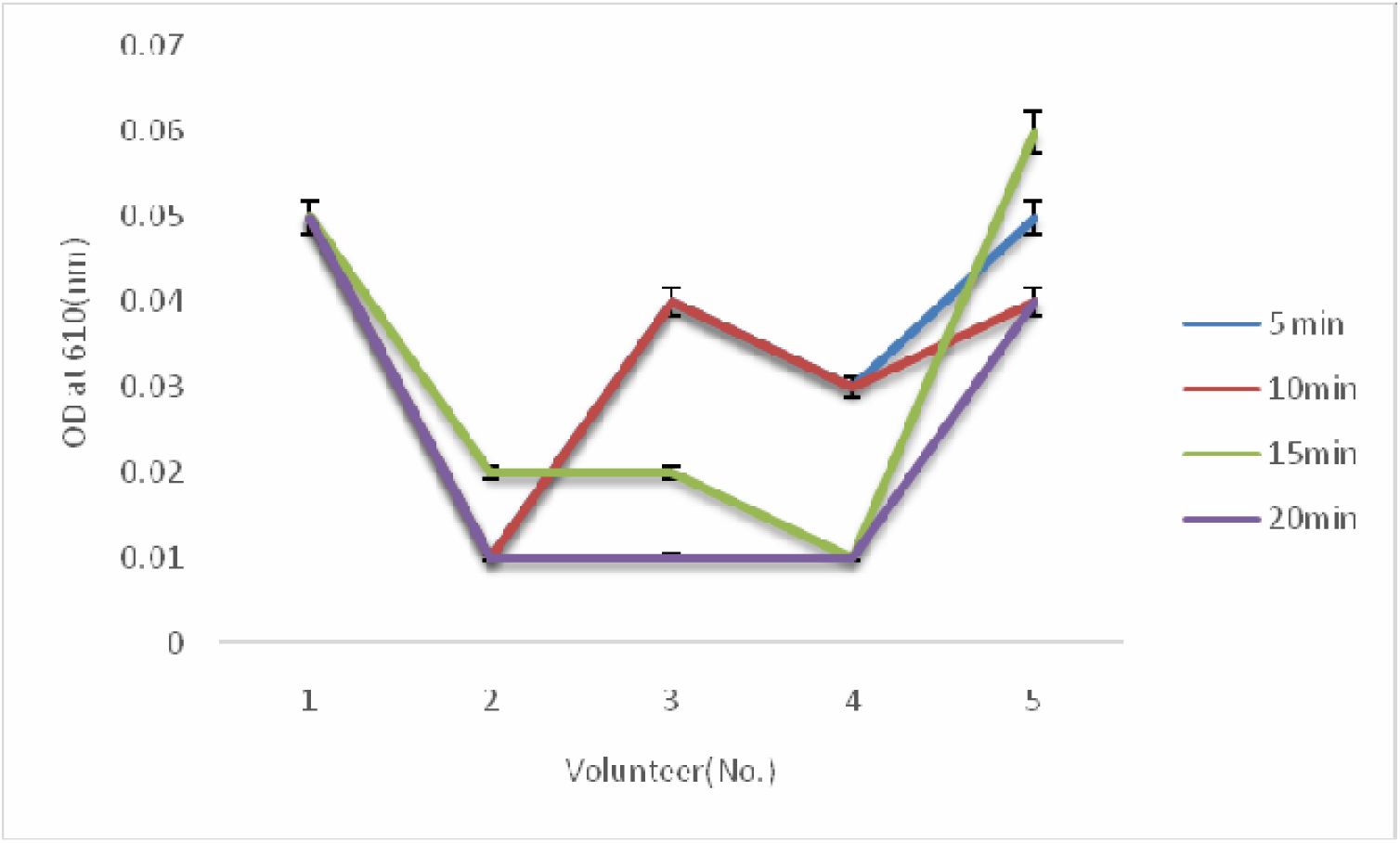
3^rd^ day observation of antimicrobial activity of Hand sanitizer against hand swab

### Antioxidant assay

#### DPPH assay (1, 1-diphenyl-2-picrylhydrazyl)

200μl of analytical sample solution and 800μl of 0.1 M Tris-HCL buffer (PH 7.4) were added into test tube or sampling tube 1ml of the DPPH Solution was added. Immediately, the solution was mixed with a test tube mixer for 10 s. Thereafter, it was left at room temperature in the dark. Exactly 30 minute after the addition of the DPPH solution, the absorbance of the solution at 517 nm was measured. A mixed solution of 1.2 mL of ethanol and 800μl of Tries-HCL buffer was used as the blank. The absorbance at the addition of the analytical sample was expressed as, the absorbance at the addition of ethanol instead of the sample as Ac, and the inhibition ratio (%) was obtained from the following equation.

Inhibition ratio (%) = (Ac-As)/Ac) □ 100

In the analytical procedure distributed, the measurement at six points of concentration, including control, was required. The measurement of the DPPH radical scavenging activity for the analytical sample solution at each concentration was repeated three times.

#### Stability

The stability studies were carried out by storing at different temperature conditions like 40□C, 25□C & 37□C for 4 weeks. During the stability studies no change in color and no phase separation were observed in the formulated hand wash. Also the formulations with stand its activity

## RESULTS AND DISCUSSION

According to the zone of inhibition formed resulting from the herbal hand wash gel against different bacterial isolates, showed that the hand wash prepared with *Azadirachta indica, Ocimum sanctum* and *Citrus limon* extracts had great activity. Statistical analysis findings in fig: 6 showed that herbal hand wash gel is the broad spectrum antibacterial agent with different response for different bacterial kinds tested. From the investigation it was clear *that Azadirachta indica, Ocimum sanctum* and *Citrus limon* were equally effective against both the groups of bacteria. *Azadirachita indica* produced the widest zone of inhibition against *E.coli* with diameter of 3.2 cm, *S.aureus* 2.8 cm, *Salmonella* 2.4 cm followed by *C.albicans* 2.6 cm. *Ocimum sanctum* produced the widest zone of inhibition against *E.coli* with diameter of 2.8 cm, *S.aureus* 2.5 cm, *Salmonella* 2.0 cm followed by *C.albicans* 2.8 cm. *Citrus limon* produced the widest zone of inhibition against with diameter of *C.albicans* 3.6 cm, *S.aureus* 3.4cm, *Salmonella* 3.2 cm followed by *E.coli* 3.0 cm. Formulated hand sanitizer produced the widest zone of inhibition against *Salmonella* 3.7 cm *E.coli* with diameter of 3.6 cm, *S.aureus* 3.2 cm, followed by *C.albicans* 3.1 cm The inhibition by *Azadirachta indica, Ocimum sanctum* and *Citrus limon* could be due to the presence of active constituents such as nimbn, nimbinin, nimbidin, nimbosterol, cirsilineol, circimaritin, isothymusin, apigenin and rosameric acid, orientin, and vicenin and Limonene. These are terpenoids in nature. Their activity is a function of the lipophilic properties of the constituent terpenes, the potency of their functional groups [11].

Hand sanitizer showed the highest anti-oxidant activity of 11.02% while *Ocimum sanctum* showed 10.76%, *Azadirachta indica* showed 6.18% and followed by *Citrus limon* showed 1.08% DPPH scavenging activity (Table 5). These phenomena can be attributed towards the presence of phenolic constituents in formulation and herbal extract which in turn indicates the stability of the formulation. Swab sample test of formulated hand sanitizer on 5 different volunteers on randomly selected 3 different days evoked an encouraging response. On day first this product reduced and holds the microbial load under significantly low concentration for 15 mins whereas for 2^nd^ day test it completely reduced the load on 20 min time period. On 3^rd^ day test it was shown some increase in microbial load behavior but that could be explain with erratic nature of exposure towards the environment. The microbial load difference between before and after of hand sanitizer use was not significant enough to indicate any contradiction towards its antimicrobial efficacy. Like cosmetics, cosmeceuticals are topically applied but they contain ingredients that influence the biological functions of skin. Traditional Indian medicine indicating the potential medicinal value of *Azadirachta indica, Ocimum sanctum* and *Citrus limon*. Natural remedies are more acceptable in the belief as they are safer with fewer side effects than the synthetic ones. Herbal formulations have emergent demand in the global market. It is an attempt made to establish the herbal gel based alcohol free natural hand sanitizer containing *Azadirachta indica, Ocimum sanctum* and *Citrus limon* extract at various concentrations.

**Table 5:** Result of antimicrobial study of DPPH scavenging activity of samples

| Sample name | O.D value | % inhibition |
| --- | --- | --- |
| Standard | 0.9797 |  |
| *Azadirachta indica* | 0.9191 | 6.18% |
| *Ocimum sanctum* | 0.8742 | 10.76% |
| *Citrus limon* | 0.9692 | 1.07% |
| Hand sanitizer | 0.8640 | 11.02% |

## CONCLUSION

Hand hygiene can also be a problem in between the people. Prevention and control of infectious activities are designed to limit the spread of infection and provide a safe environment for all people, regardless of the setting. In light of the emergence of antibiotic resistant organisms, effective infection control measures, such as hand sanitizing, are essential to prevention. Hand sanitizer gels are used for the purpose of cleaning hands. Its composition is prepared according to delicateness of skin so that it cannot cause any type of irritation.

It is concluded that from the result that the gel formulation is good in appearance, homogeneity. This preliminary *in vitro* study demonstrated that *Azadirachta indica, Ocimum sanctum* and *Citrus limon* extracts active against hand swab sample. Natural gel based alcohol free hand sanitizer was as effective against pathogenic bacteria in volunteer’s samples with no side effects on human tissue. Thus these compounds can be extracted and incorporated in hand wash formulation in order to prepare superior antiseptic herbal hand wash gel with little or no side effects. Thus, a new way can be found to provide safe and healthier living through germ-free hands. Although the removal is not 100% but a major number can be reduced.

## Acknowledgement

The authors express their gratitude to Patanjali Groups, for the moral support throughout the research project.

